# Landscape of Differentiation Potentials as a “Hallmark” in Oral-derived MSCs

**DOI:** 10.1101/2024.08.02.606413

**Authors:** H Chopra, C Cao, H Alice, S Kak, B Maska, R Tagett, J Sugai, L Garmire, D Kaigler

**Affiliations:** Kaigler lab of stem cell science and Regenerative Medicine, Department of Periodontics and Oral Medicine, School of Dentistry, University of Michigan, Ann Arbor, MI-48109, USA; Bioinformatics Core, University of Michigan, Ann Arbor, MI-48109, USA; Computational Medicine & Bioinformatics, University of Michigan, Ann Arbor, MI-48109, USA; Department of Biomedical Engineering, University of Michigan; Ann Arbor, MI-48109, USA

## Abstract

**Background:** Mesenchymal stem cells (MSCs) offer clinical promise for use in cell therapy approaches for regenerative medicine. A therapeutic challenge is that MSCs from different tissues are phenotypically and functionally distinct. Therefore, this study aims to molecularly characterize oral-derived MSCs by defining one of the three hallmarks of MSCs, differentiation potential, to discern their true molecular identities.

**Methods:** Three different populations of oral tissue MSCs (from alveolar bone-aBMSCs; from dental pulp-DPSCs; and from gingiva-GMSCs) from three different patients were isolated and cultured. These MSCs were characterized for their stemness by flow cytometry and multi-differentiation potential, and their RNA was also isolated and analyzed quantitatively with RNA sequencing. Total mRNA-seq was performed and differentially expressed genes (DEGs) were identified in pairwise (DPSCs vs. aBMSCs, GMSCs vs. aBMSCs, and GMSCs vs. DPSCs) and tissue-specific comparisons (aBMSCs vs. Others, DPSCs vs. Others, GMSCs vs. Others) (FDR, *p<0.05*). Further, these DEGs, either common between MSC populations or unique to a specific MSC population, were evaluated for pathways and biological processes

**Results:** aBMSCs, DPSCs, and GMSCs were successfully isolated and characterized. The tissue-specific comparison revealed that DEGs were most numerous in DPSCs (693 genes) as compared to aBMSCs (103 genes) or DPSCs (232 genes). Statistically significant DEGs through pairwise comparisons present higher numbers in GMSCs vs. DPSCs (627) as compared to either DPSCs vs aBMSCs (286) or GMSCs vs. aBMSCs (82). Further analysis found that RUNX2, IBSP, SOX6, ACAN, and VCAM1 were significantly upregulated in aBMSCs. In DPSCs, BMP4 and IL6 were significantly downregulated, whereas AXL and NES were significantly upregulated. In GMSCs, AGPT1, SEMA4D, and PGDFA were significantly downregulated. Additionally, MAPK, PI3-AKT, and RAS signaling pathways were significantly regulated in GMSCs. Interestingly, aBMSCs and DPSCs revealed positive regulation of osteoblast differentiation, whereas GMSCs revealed negative regulation of osteoblast differentiation. DPSCs also revealed negative regulation of angiogenesis.

**Conclusions:** Oral-derived MSCs have an inherent “landscape” of differentiation defined by their tissue of origin; yet this differentiation potential can be modulated by their microenvironment.

**Graphical Abstract:** 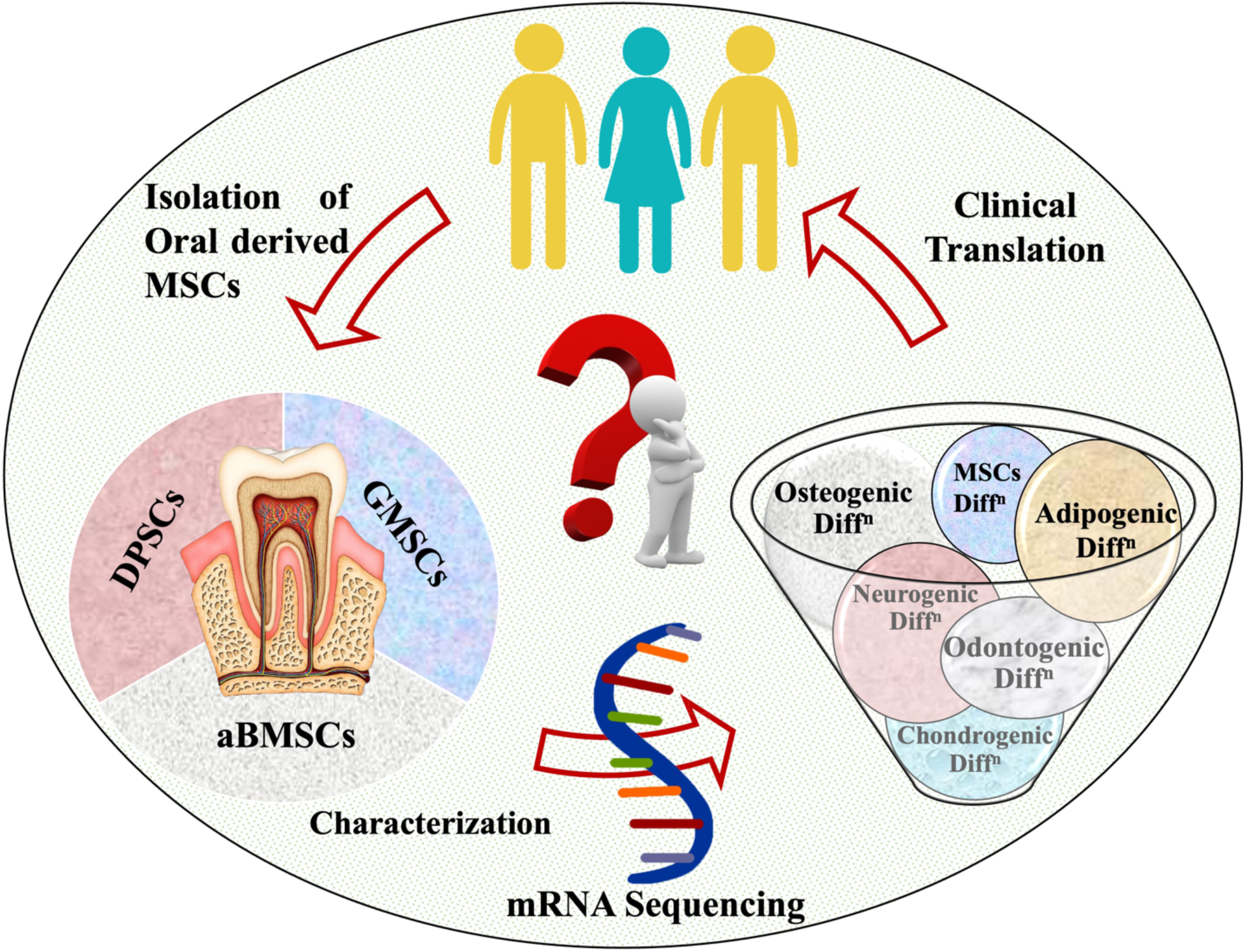

## Background

Mesenchymal stem/stromal cells (MSCs) have garnered significant attention in regenerative medicine because, besides their tissue repair and regenerative capacity, they are relatively easy to isolate, have immunomodulatory properties [1], have multilineage differentiation potential [2], and are associated with significantly fewer ethical concerns regarding their use vs. embryonic stem cells. In recent years, MSCs have been isolated from various dental and non-dental tissues [1, 3]. Correspondingly, there has been a significant increase in MSC-related clinical trials from one trial in 1995 to 13,300 studies in 2023 (ClinicalTrials.gov), underlying their potential efficacy in treating various clinical conditions [4, 5].

The heightened interest in their use underscores the ambiguities and inconsistencies in defining what a “true” MSC is. To address these issues, criteria to characterize these MSCs have been established as 1) adherence to plastic, 2) expression of specific surface antigens, and 3) multilineage differentiation potential [6]. However, it is increasingly recognized that MSCs derived from distinct tissue sources exhibit notable heterogeneity in their phenotypic and functional properties [7, 8]. Further, the heterogeneity of MSCs may be due to various biological factors, such as age, gender, health condition, and genetic background, as well as technical factors, such as the different strategies to isolate and expand MSCs [9–11]. This divergence highlights the importance of comprehensively characterizing MSCs at the molecular level to better understand their regenerative potential and most appropriate therapeutic uses.

Although flow cytometry and fluorescence-activated cell sorting (FACS) have been the status quo in the identification and isolation of MSC subsets by detecting the surface markers that define these MSCs for the last 20 years [6, 12], these techniques fail to pinpoint molecular signatures that are characteristic of different MSC populations. Similarly, other molecular methods, such as magnetic bead assay, can identify and sort functionally diverse MSC population subsets, but the molecular nature of many MSCs/subsets remains unclear. In this aspect, applying transcriptomics for MSC characterization can provide important insights into their molecular heterogeneity and underlying disposition for differentiation.

Over the past decade, RNA-seq has become an indispensable tool in molecular biology for transcriptome-wide analysis of differential gene expression and understanding overall pathway activity across cell types and different conditions [13, 14]. As such, this tool lends itself to being used to characterize the molecular heterogeneity of MSCs rigorously. In doing so, these insights would enable us to better understand mechanisms underlying the phenotypic and functional differences seen between different MSCs.

Oral-derived MSCs represent unique MSC populations in that they can be derived from different tissues within the same macroenvironment (i.e., oral cavity) but from different microenvironments (i.e., bone, gingiva, tooth). These different microenvironments create distinct niches, which are thought to contribute to the differences reported in how these MSC populations behave relative to one another. The phenotypic differences in oral-derived MSCs are primarily responsible for conflicting reports about their differentiation capacity, particularly with gingiva-derived MSCs (GMSCs) and dental pulp-derived MSCs (DPSCs) [15, 16]. Further, as alveolar bone-derived MSCs (aBMSCs) have been recently discovered as compared to other oral-derived stem cells [17], there is much more to understand about these MSCs. Therefore, in this study, we aim to molecularly characterize aBMSCs, DPSCs, and GMSCs by bulk RNA-seq. to better define their molecular properties. By evaluating the molecular biology “landscape” of these MSCs across various differentiation potentials, we aim to unravel the intrinsic identity of these MSCs to elucidate the molecular mechanisms underpinning their unique characteristics.

## Materials and Methods

### Human Tissue Sampling

Human tooth, alveolar bone, and gingiva from different patients (n=3) undergoing routine oral surgical procedures requiring tooth removal were obtained from the Dental Clinics at the School of Dentistry, The University of Michigan, with approval from the Institutional Review Board at The University of Michigan (IRB #HUM00142680).

### Isolation and culturing of oral-derived mesenchymal Stem cells

After transferring the tissues to a sterile container containing α modification of minimum essential media (α-MEM, Thermo Fisher Scientific, Waltham, MA, USA) and 1% antibiotic-antimycotic (Thermo Fisher Scientific), the tissues were rinsed thoroughly with phosphate-buffered saline (PBS). Alveolar bone mesenchymal stem cells (aBMSCs), dental pulp stem cells (DPSCs), and gingival mesenchymal stem cells (GMSCs) were isolated from the alveolar bone, dental pulp of the tooth, and gingiva, respectively from three different patients according to the previously described procedure [18–20].

In brief, MSCs from all the tissues except aBMSCs were isolated by enzymatic digestion method using collagenase/dispase (Worthington Biochemical, Lakewood, NJ, USA) [18–20]. The fragments of alveolar bone and cell suspension from the dental pulp, or gingiva, were placed in a T-25 flask with their respective growth medium and maintained at 37°C with 5% CO_2_ in a humidified atmosphere, with the medium replaced every three days. For aBMSCs and DPSCs, nucleosides-containing α-MEM (Thermo Fisher Scientific) was supplemented with 15% fetal bovine serum (FBS; MilliporeSigma, Burlington, MA, USA), 0.1 mM L-ascorbic acid-2-phosphate (AA2P; MilliporeSigma), and 1% antibiotic-antimycotic, whereas for GMSCs, the composition of the media remained the same as that used with aBMSCs or DPSCs except with the addition of non-essential amino acids (1%, Thermo Fisher Scientific) and reduced concentration of FBS to 10%. After colonies of adherent cells appeared in one to two weeks, at 70% confluency, these adherent cells were passaged and cultured in their respective growth medium. All the aBMSCs, GMSCs, and DPSCs used in the present study were used from the passage (P) P3-P5, with the same passage used consistently within each experiment.

### Multipotent differentiation potential

aBMSCs, GMSCs, and DPSCs cells were evaluated for their multidifferential potential by osteogenic, adipogenic, and neurogenic differentiation. Further, each experiment was replicated thrice.

#### Osteogenic differentiation

aBMSCs, GMSCs, and DPSCs at P4 were seeded in 6-well plates at a density of 1.5 × 10^4^ cells/well in their respective culture medium for 24 h at 37 °C in a humidified atmosphere containing 5% CO_2_. After 24 h, the medium in the treatment group was changed to customized osteogenic differentiation media containing 85% α-MEM, 10% FBS, 1% antibiotic-antimycotic, 2% Glutamax (Thermo Fisher Scientific), 0.5% (50 µg/mL) L-ascorbic acid-2-phosphate, 0.5% (10 nM) dexamethasone, and 1% (10 mM) β-glycerophosphate (All from MilliporeSigma). On the other hand, the control group was supplemented with only α-MEM, 15% FBS, 1% antibiotic-antimycotic, and 2% Glutamax. Both the mediums were refreshed at 3-day intervals for four weeks. The wells were then evaluated for osteogenic differentiation by histochemical staining (HS) with Alizarin Red S (ARS) (pH:4.1–4.3; MilliporeSigma). After HS, the plates were then scanned with a scanner (Hewlett Packard, USA)

#### Adipogenic differentiation

aBMSCs, GMSCs, and DPSCs at P4 were seeded at a density of 2× 10^4^ cells in 6-well plates and incubated at 37 °C in a humidified atmosphere of 5% CO_2_. Their respective culture medium was replaced every three days until the cells reached 100% confluence (3–4 days). At near 100% confluence, α-MEM was removed, and three cycles of adipogenic induction media (AIM) and adipogenic maintenance media (AMM) were initiated in treatment groups. In comparison, the control group was supplemented with only 87% α-MEM, 15% FBS, 1% antibiotic-antimycotic, and 2% Glutamax. The AIM was composed of α-MEM, 15%, FBS 1% antibiotic-antimycotic, 2% Glutamax, 10mM 3-isobuty-l-methyl-xanthine, 100 µM dexamethasone, 50 mM indomethacin, and 0.1% h-insulin (recombinant), whereas AMM was composed of α-MEM, 15% FBS, 1% antibiotic-antimycotic, 2% Glutamax, and 0.1% h-insulin (recombinant) (All from MilliporeSigma). After three cycles, both the groups were maintained in AMM for seven days with the replacement of AMM twice at an interval of 3 days. The wells were then analyzed for adipogenic differentiation by HS with Oil Red O (ORO) (MilliporeSigma).

#### Neurogenic differentiation

aBMSCs, GMSCs, and DPSCs at P4 were seeded in a 6-well plate at a density of 1.5 × 10^4^ cells/well and incubated in the culture medium for 24 h at 37 °C in a humidified atmosphere containing 5% CO_2_. After 24 h, the medium in the treatment group was changed to customized neurogenic differentiation media containing Neurobasal-A medium supplemented with 2% B-27 (Both from Gibco-Lifesciences) with 20 ng/mL epidermal growth factor, 40 ng/mL fibroblast growth factor (Both from BD Biosciences, San Jose, CA, USA), 1% antibiotic-antimycotic and 2% Glutamax. The control group was supplemented with α-MEM, 1% antibiotic-antimycotic, and 2% Glutamax. Both the mediums were refreshed at 3-day intervals for three weeks. The wells were then analyzed for neurogenic differentiation with Cresyl Violet (pH:3.7–3.9; MilliporeSigma).

### Histochemical staining (HS)

For the osteogenic assay, after four weeks of culture, mediums of both the treatment and control groups were removed. The wells were then washed twice with PBS for 2 min, followed by incubation of cells with 4% paraformaldehyde (PFA) for 30 min at room temperature (RT). After fixation, PFA was removed, and the wells were again washed twice with PBS for 5 min. Freshly prepared ARS was added to all wells and incubated for 45 min at RT. After staining, ARS was removed and washed with PBS until the excess stain was no longer apparent. The plates were then scanned with a scanner, and images of the cells were taken using the phase contrast microscope.

The same protocol was followed for the neurogenic assay, except that a freshly prepared Cresyl Violet stain was used.

For the adipogenic assay, a working solution of ORO was prepared by mixing three parts (30 mL) of ORO stock solution with two parts (20 mL) of deionized water, followed by filtering with a 0.2 µm syringe filter. All the remaining procedures were the same as described above, except that after fixation, 60% isopropanol (MilliporeSigma) was added to each well for 5 min at RT, followed by staining with ORO.

### Flow cytometry

Flow cytometry was used for quantitative analysis of the phenotype of the cells as described by Dominici et al. [6]. aBMSCs, GMSCs, and DPSCs from all three patients at P4 were cultured in a T75 flask and after reaching 70% confluency, the monolayers were trypsinized, filtered and samples were prepared at a concentration of 0.2 × 10^6^ to 0.5 × 10^6^ cells. The cells were simultaneously blocked with 0.1% bovine serum albumin (BSA) and incubated with the conjugated antibody (Table S1) for 30–45 min in an ice box with gentle stirring. After incubation, the samples were washed thrice with PBS. The negative control consisted of unstained cells, whereas the isotype control had cells with the isotype of the corresponding antibody and incubated for 30–45 min, followed by washing thrice for 5 min in PBS. All the samples were strained with a 70 µm filter to obtain singlets and then subjected to Biorad ZE5 (Biorad, California. USA) to analyze respective markers. A minimum of 10,000 events were recorded.

### RNA extraction and quality control

aBMSCs, GMSCs, and DPSCs at P4 from all three patients were cultured in T25 flasks. At 70% confluency, the monolayers were briefly rinsed with PBS and then lyzed with RLT Lysis Buffer from the RNeasy Mini Plus Kit or RNeasy Micro Plus Kit (Qiagen, Hilden, Germany) with the addition of 1% 2-mercaptoethanol. The lysates were kept at −80 °C until processed to extract RNA with the rest of the reagents from the kits according to the manufacturer’s instructions. The RNA concentrations were determined by Qubit Assay using the Qubit RNA HS Assay Kit and the Qubit fluorometer (Thermo Fisher Scientific) according to the manufacturer’s instructions. The quality of RNA was examined using a Bioanalyzer (Agilent 2100, Agilent Technologies).

### mRNA enrichment and library preparation

Poly-A Selection was used to enrich mRNA using oligo (dT) beads that bind to the poly-A tail of mRNA molecules. After enrichment, double-stranded cDNA was synthesized, followed by fragmentation and ligation of sequencing adapters to the ends of the cDNA fragments using a Lexogen QuantSeq library preparation kit. The adapter-ligated cDNA fragments were amplified using polymerase chain reaction (PCR) to generate sufficient material for sequencing.

### RNA sequencing and analysis

RNA sequencing was performed by the University of Michigan Advanced Genomic Core using Illumina. The quality of the raw sequencing data was assessed using FastQC [21] followed by Cutadapt [22] to find and remove adapter sequences, primers, poly-A-tails, and other types of unwanted sequences in the high-throughput sequencing reads. FastQC was repeated on the trimmed reads. The trimmed reads were aligned to the Homo sapiens reference genome (GRCh38.p14 - hg19) using STAR [23] and data was normalized using the trimmed mean of M values (TMM) technique [24]. Differential gene expression analysis was performed with DESeq2 [25], using a negative binomial generalized linear model in edge R [26] and accounting for natural variation of patients by using a pair-sample regression design.

### Data analysis

The flow cytometer data was analyzed using FlowJo Version 10.0 (FlowJo, LLC, Ashland, OR, USA), and values are presented as mean. For analysis of differential gene expression, *p-values* were adjusted for multiple comparisons using the Benjamini-Hochberg false discovery rate (FDR) [27]. Comparing two MSC types, significant differential expression was determined based on FDR of ≤0.05 and linear fold-change ≥1.5 was reported as significantly differentially expressed (DE). These DE genes (DEGs) were analyzed as pairwise comparisons (DPSCs vs. aBMSCs, GMSCs vs. aBMSCs, and GMSCs vs. DPSCs). Tissue-specific comparisons were also made, comparing one MSC type to the two others based on their fold-change and FDR statistics. If a given MSC type was significantly differentially expressed in both pairwise comparisons in which it was included, and if the fold change concerning the two other MSC types was in the same direction, then that given MSC was considered differentially expressed versus the two others. For significant DE genes in the tissue-specific comparisons (aBMSCs vs. Others, DPSCs vs. Others, and GMSCs vs. Others), fold changes were assigned by averaging the fold changes in the pairwise comparisons, and FDR values were assigned using the geometric mean. For non-significant genes, fold-changes were assigned in the same way, but FDR values were set to one to avoid cases where only one of the pairwise comparisons was significant, but its FDR was significant enough to cause the combined FDR to be significant also. All these results (pairwise and tissue-specific) were fed into the iPathwayGuide (AdvaitaBio Corporation, https://ipathwayguide.advaitabio.com/), a web-based functional analysis tool that provides pathway and enrichment analyses and meta-analyses of multiple comparisons. Further, each comparison was evaluated in terms of pathways and Biological Processes from the Gene Ontology (GO). Some notable genes (Table S2, Additional File 1) and 21 GO Biological Processes. (GOBPs) were selected [28] (Table S3, Additional File 1) out of a total of 26,994 GOBPs[29] to explain the landscape of differentiation potential in all three MSC populations.

## Results

### Human tissue sampling and isolation and culturing of oral-derived MSCs

Normal human dental tissue, such as teeth, alveolar bone, and gingiva were successfully harvested from three patients without any caries, inflammation, or malignancy. aBMSCs, GMSCs, and DPSCs from these tissues for all three patients were isolated successfully and cryopreserved at P2 for subsequent investigation (Fig. S1, Additional File 1).

### Multipotent differentiation potential

aBMSCs, GMSCs, and DPSCs from all three patients differentiated into osteoblast-like cells/adipocyte-like cells/neuron-like cells in osteogenic/adipogenic/neurogenic medium producing calcific nodules (Fig. 1B)/lipid vacuoles (Fig. 1C)/nissl bodies (Fig. 1D)/ that appeared red on staining with ARS/orange on staining with ORO/blue on staining with cresyl violet, respectively. A representation of osteogenic differentiation, adipogenic differentiation, and neurogenic differentiation in aBMSCs, DPSCs, and GMSCs from all three patients is shown in Fig. 1.

**Figure 1.**
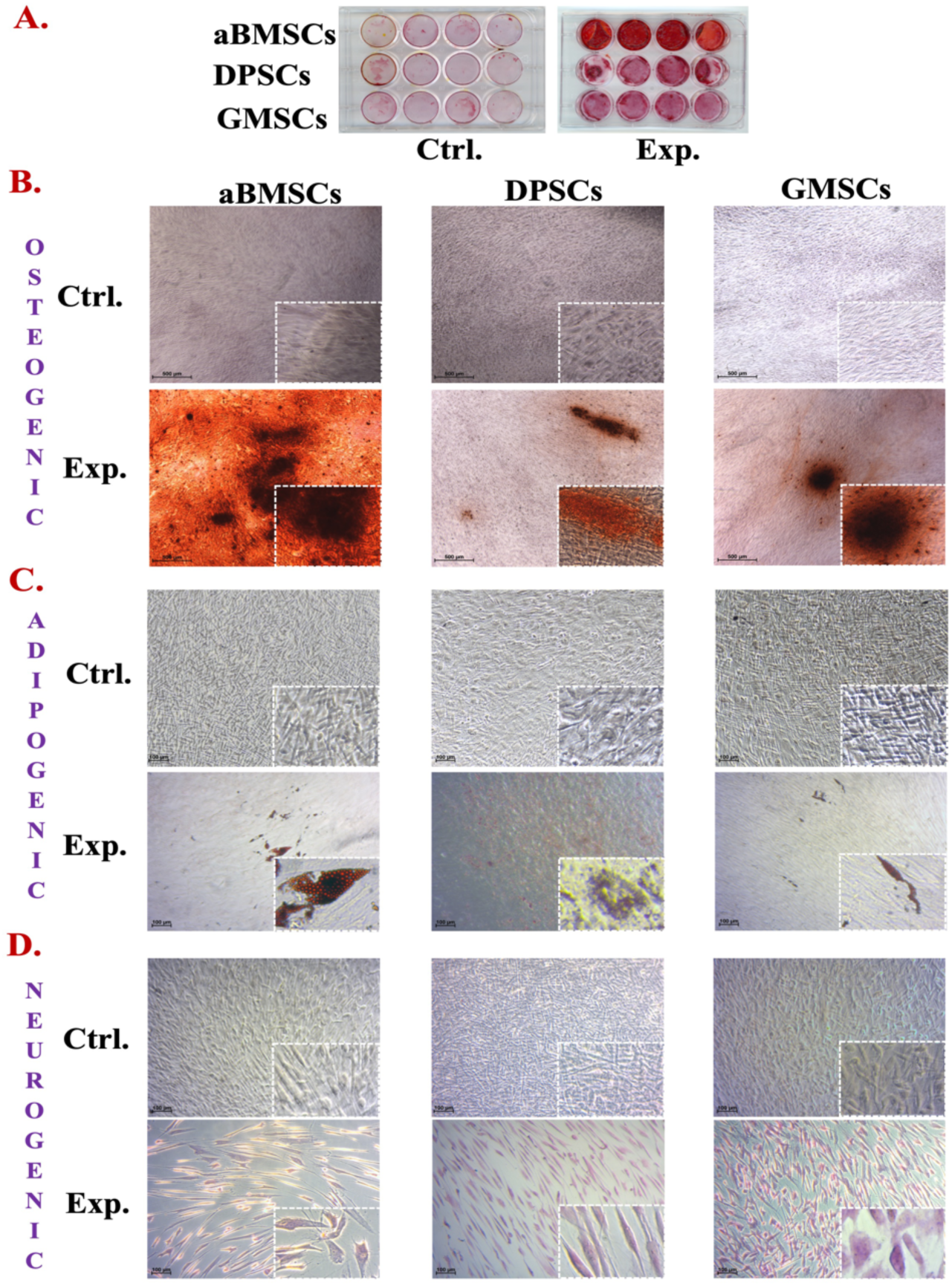
Multidifferentiation potential of aBMSCs, DPSCs, and GMSCs. The MSCs were cultured in osteogenic (4 weeks), adipogenic (3 weeks), and neurogenic (3 weeks) induction media and stained with alizarin red stain, oil red o, and cresyl violet. **A.** Visual examination of the representative osteogenic plate after osteogenic induction for four weeks. **B.** Microscopic examination (4X) of representative osteogenic plate after osteogenic induction for four weeks showing calcific nodules (10X). **C.** Microscopic examination (10X) of the representative adipogenic plate after the adipogenic induction for three weeks showing lipid vacuoles (40X). D. Microscopic examination (10X) of the representative neurogenic plate after the neurogenic induction for three weeks showing Nissl bodies at (40X).

### Flow Cytometry

Flow cytometry confirmed the characteristic phenotype of MSCs for aBMSCs, GMSCs, and DPSCs from all three patients with CD90, CD105, and CD73 showing greater than 95% expression, whereas CD45 and CD34 demonstrated less than 5% expression. An average expression of CD90, CD105, CD73, CD45, and CD34 with representative histograms from one of the patients was shown in Fig. 2.

**Figure 2.**
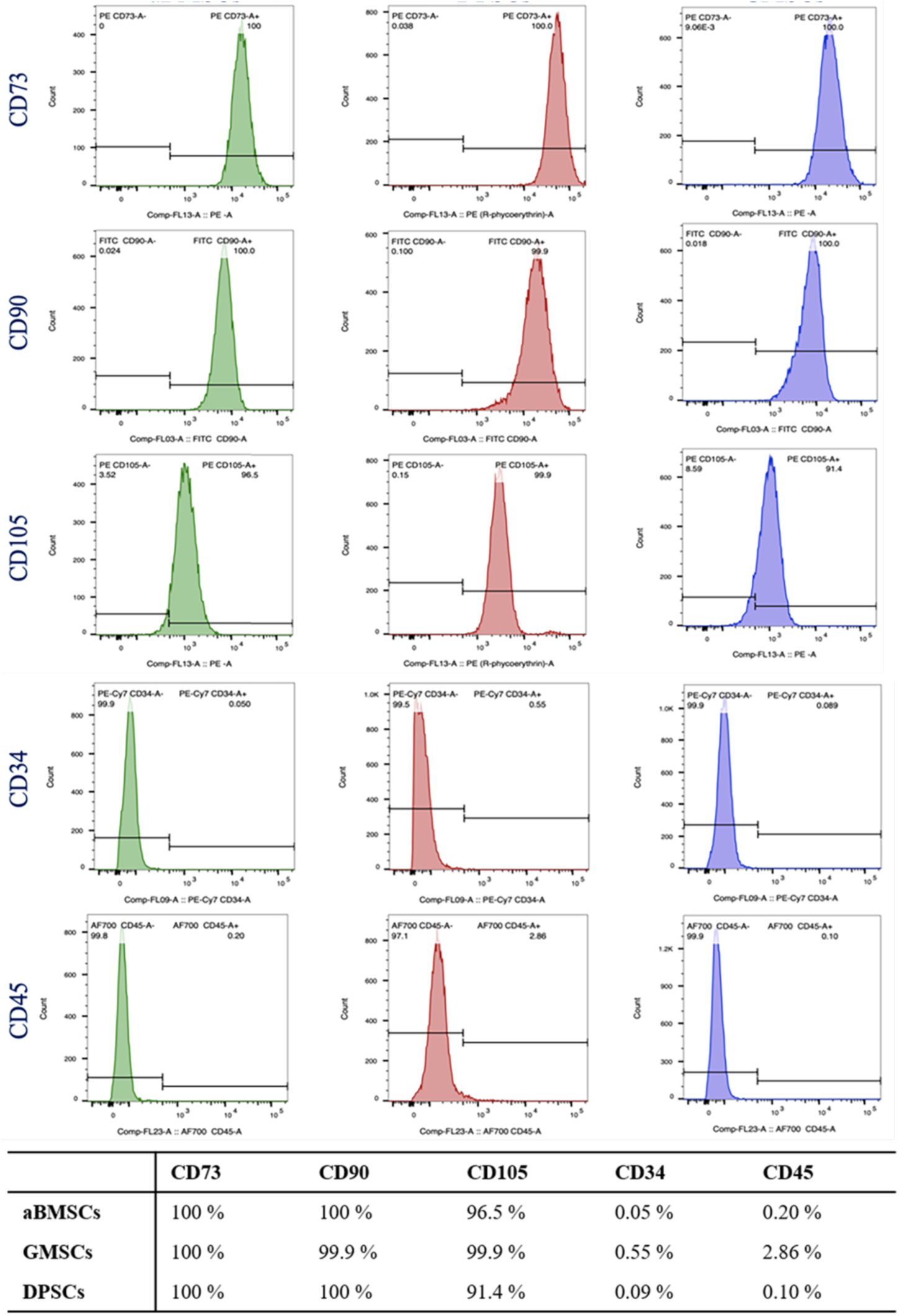
Phenotypic Characterization Using Flow Cytometry. **A.** Representative expression of MSC surface markers in aBMSCs, DPSCs, and GMSCs. **B.** The expressions of CD73, CD90, and CD105 in percentage from one of the representative patients. All three patients fulfilled the Dominici et al [6] criteria with CD105, CD73, and CD90 expressed ≥95%, whereas CD45 and CD34 were expressed ≤2%.

### RNA extraction and quality and quantity of RNA

RNA was successfully extracted from aBMSCs, GMSCs, and DPSCs from all three patients. Qualitative analysis of all nine samples revealed RIN >9 without DNA contamination.

### Pairwise comparison

#### Some genes are common to the MSC population

After a meta-analysis of pairwise differential expression comparisons, it was found that unique statistically significant DEGs were found to be higher in GMSCs vs. DPSCs (627) as compared to either DPSCs vs. aBMSCs (286) or GMSCs vs. aBMSCs (82) (Fig 3A). BMP2

**Figure 3.**
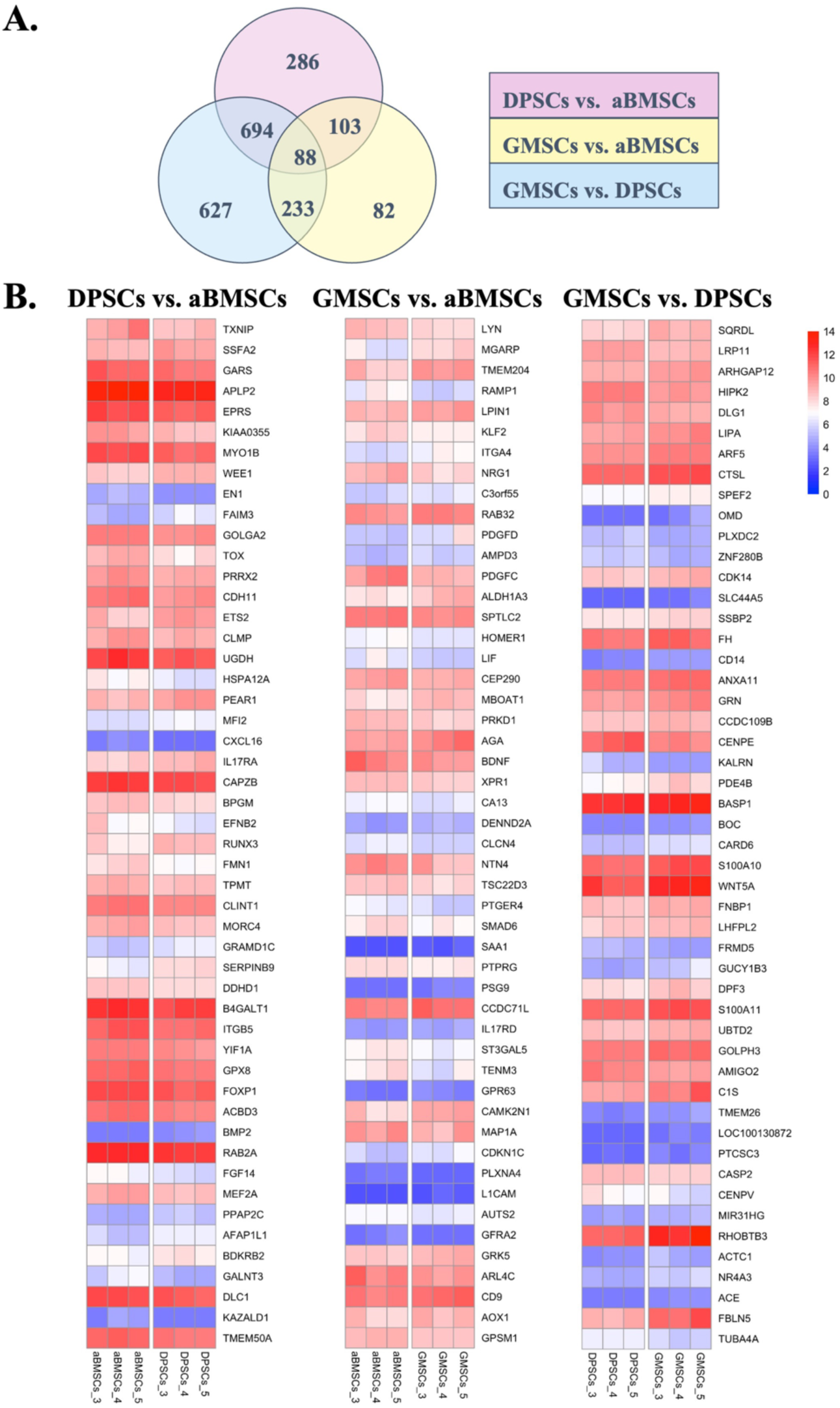
Venn Diagram and Heat Maps of Pairwise Comparisons. **A.** Venn diagram obtained by metanalysis of all DEGs in DPSCs vs. aBMSCs, GMSCs vs. aBMSCs, and GMSCs vs. DPSCs **B.** Heat maps of three different oral-derived MSC populations in a pairwise comparison showing the top 50 genes in DPSCs vs. aBMSCs, GMSCs vs. aBMSCs, and GMSCs vs. DPSCs according to their significance (FDR, *p*<0.05).

#### Some pathways are unique to two MSC populations while other pathways are common to all comparisons of MSC populations

The top 10 pathways with their corresponding genes for each pairwise comparison; DPSCs vs. aBMSCs, GMSCs vs. aBMSCs, and GMSCs vs. DPSCs after p-value correction with FDR (*p<0.05*), are shown in Fig. 5. After a meta-analysis of these comparisons, it was found that eight pathways [out of total 33 statistically significant (FDR, *p<0.05*) pathways] were common to all comparisons (Table S4, Additional file 1). Notably, the PI3-AKT signaling pathway was found to be statistically significant (FDR, *p<0.05*) in GMSCs vs. aBMSCs and GMSCs vs. DPSCs, TGF-β signaling pathway was unique and statistically significant (FDR, *p<0.05*) in GMSCs vs. DPSCs (FDR, *p<0.05*), whereas RAP1 signaling pathway, RAS1 signaling pathway, IL7 signaling pathway, and calcium signaling pathway were unique and statistically significant (FDR, *p<0.05*) in GMSCs vs aBMSCs (Table S4, Additional File 1).

#### GO Biological processes

It was found that only 1606/26,994 GOBPs were found to be statistically significant (FDR, *p<0.05*) in all pairwise comparisons. Of these 1606 Biological processes, 15/21 preselected biological processes were found with seven biological processes common to all comparisons (GO:0001649, GO:0045669, GO:0002062, GO:0032332, GO:0030182, GO:0001525, and GO:0045766). Four GOBPs were common to both DPSCs vs. aBMSCs and GMSCs vs. DPSCs (GO:0045444, GO:0045599, [GO:0016525](https://www.ebi.ac.uk/QuickGO/term/GO:0045599), and GO:0045666), one was common to both GMSCs vs. aBMSCs and GMSCs vs. DPSCs ([GO:0045668](https://www.ebi.ac.uk/QuickGO/term/GO:0045666)), whereas two GOBPs were unique to GMSCs vs. DPSCs (GO:0045665 and GO:0032331), and one GOBP was unique to GMSCs vs. aBMSCs (GO:2000741) (Table 1).

**Table 1:**
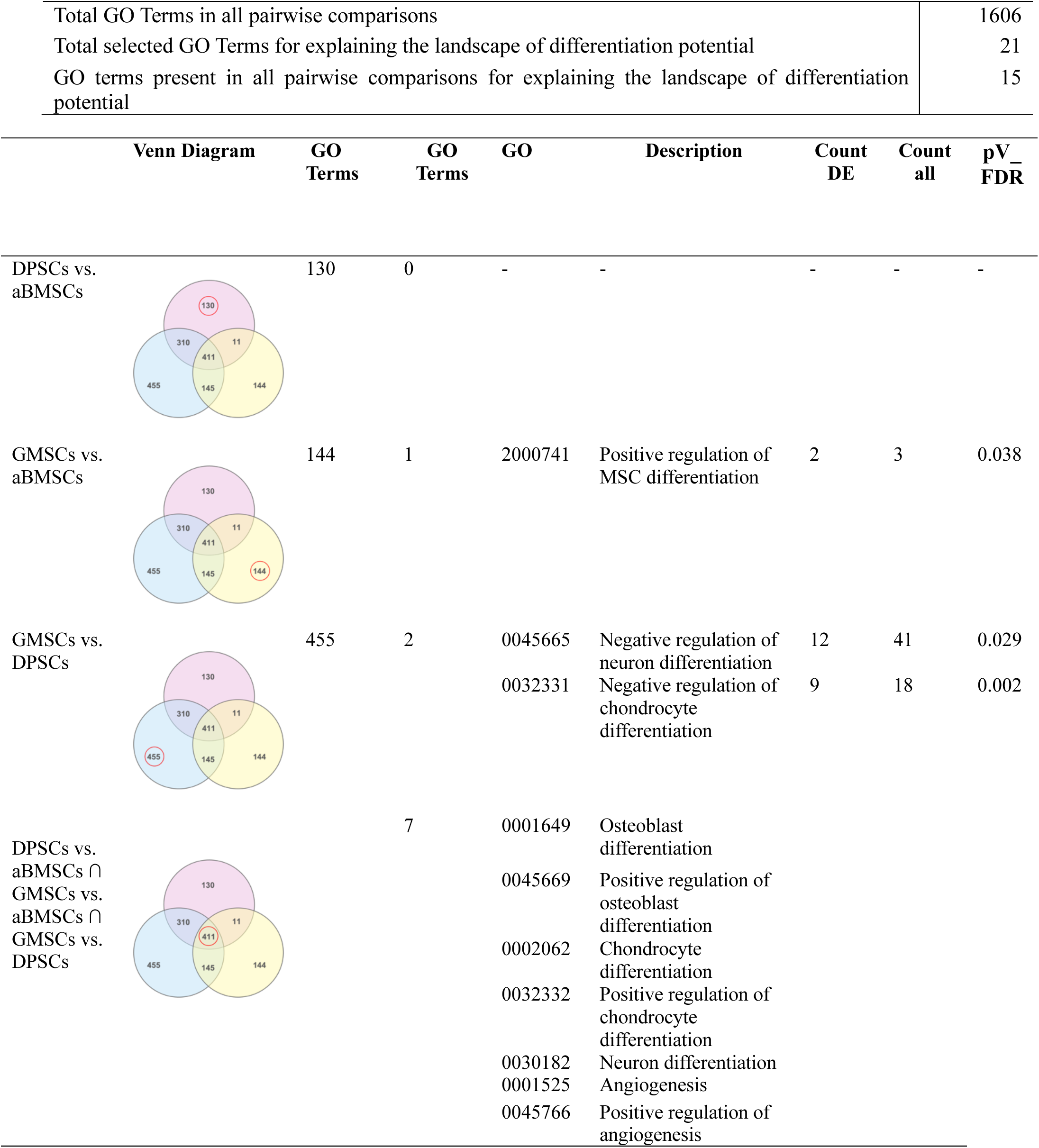
Pairwise comparison of three MSC populations showing unique GO terms, DEseq genes, and significance of these GO Terms uniquely to their comparisons (p-value correction: FDR)

| Total GO Terms in all pairwise comparisons |  |  |  |  |  |  | 1606 |
| --- | --- | --- | --- | --- | --- | --- | --- |
| Total selected GO Terms for explaining the landscape of differentiation potential |  |  |  |  |  |  | 21 |
| GO terms present in all pairwise comparisons for explaining the landscape of differentiation potential |  |  |  |  |  |  | 15 |
| Venn Diagram | GO Terms | GO Terms | GO | Description | Count DE | Count all | pV_FDR |
| DPSCs vs. aBMSCs | 130 | 0 | - | - | - | - | - |
| GMSCs vs. aBMSCs | 144 | 1 | 2000741 | Positive regulation of MSC differentiation | 2 | 3 | 0.038 |
| GMSCs vs. DPSCs | 455 | 2 | 0045665<br>0032331 | Negative regulation of neuron differentiation<br>Negative regulation of chondrocyte differentiation | 12<br>9 | 41<br>18 | 0.029<br>0.002 |
| DPSCs vs. aBMSCs ∩<br>GMSCs vs. aBMSCs ∩<br>GMSCs vs. DPSCs | | 7 | 0001649<br>0045669<br>0002062<br>0032332<br>0030182<br>0001525<br>0045766 | Osteoblast differentiation<br>Positive regulation of osteoblast differentiation<br>Chondrocyte differentiation<br>Positive regulation of chondrocyte differentiation<br>Neuron differentiation<br>Angiogenesis<br>Positive regulation of angiogenesis | | | |

### Tissue-specific comparison

#### Each population of MSC is unique with its characteristic genes

It was found that DE genes in DPSCs were the most numerous (693 genes) as compared to aBMSCs (103 genes) or GMSCs (232 genes) (Figure 6A). Few of the important DE genes such as RUNX2 and IBSP (osteoblast differentiation), SMAD7, SOX6 and ACAN (chondrocyte differentiation), and VCAM1 (inflammation) were significantly upregulated in aBMSCs as compared to DPSCs or GMSCs (Fig. S2D, Additional File 1). BMP4 (osteoblast/fat cell/chondrocyte/neuron differentiation), IL6 (inflammation), PPARG (fat cell differentiation), TWIST1 (MSC differentiation), PAX9 (odontoblast differentiation), and SEMA3A (neuron differentiation) were significantly downregulated in DPSCs, whereas AXL (chondrocyte differentiation) and NES (neuron differentiation) were significantly upregulated in DPSCs as compared to aBMSCs or GMSCs (Fig. S2E, Additional File 1). In GMSCs, SEMA4D (positive regulation of phosphatidylinositol 3-kinase signaling, neuron differentiation, and regulation of phosphate metabolic process) and TWIST2 (MSC differentiation) were significantly upregulated, whereas GDNF (neuron differentiation), ANGPT1, and PDGFA (endothelial differentiation), were significantly downregulated in comparison to the other two MSCs (Fig. S2F, Additional File 1). Heat maps of the top 50 DEGs and volcano plots of specific DEGs as explained above in tissue-specific comparisons were shown in Fig. 6B and Fig. 4B, respectively.

**Figure 4.**
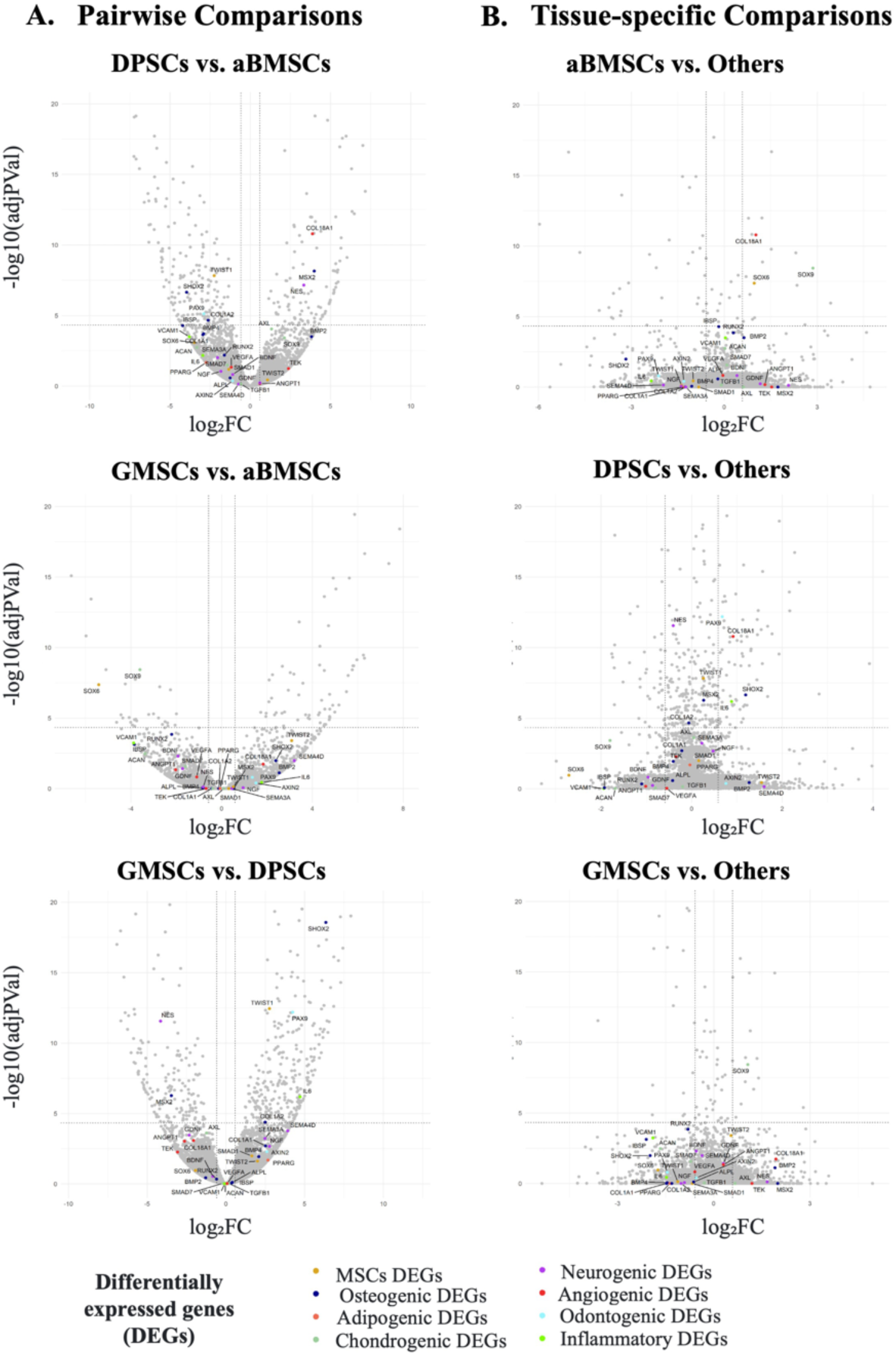
Volcano Plots of DEGs. Volcano plots of three different oral-derived MSC populations with significant DEGs were represented in terms of their measured expression change (x-axis) and the significance of the change (y-axis). The significance is described in terms of the negative log (base 10) of the p-value so that more significant genes are plotted higher on the y-axis. Further color coding and annotation of some specific DEGs related to seven differentiation activities in both comparisons were depicted. **A.** Pairwise comparison (a) DPSCs vs. aBMSCs revealed 1171 DEGs (b) GMSCs vs. aBMSCs revealed 506 DEGs (c) GMSCs vs. DPSCs revealed 1642 DEGs. **B.** Tissue-specific comparison. (a) aBMSCs vs. others revealed a significant 128 DEGs (b) DPSCs vs. others revealed a significant 771 DEGs (c) GMSCs vs. others revealed a significant 305 DEGs.

**Figure 5.**
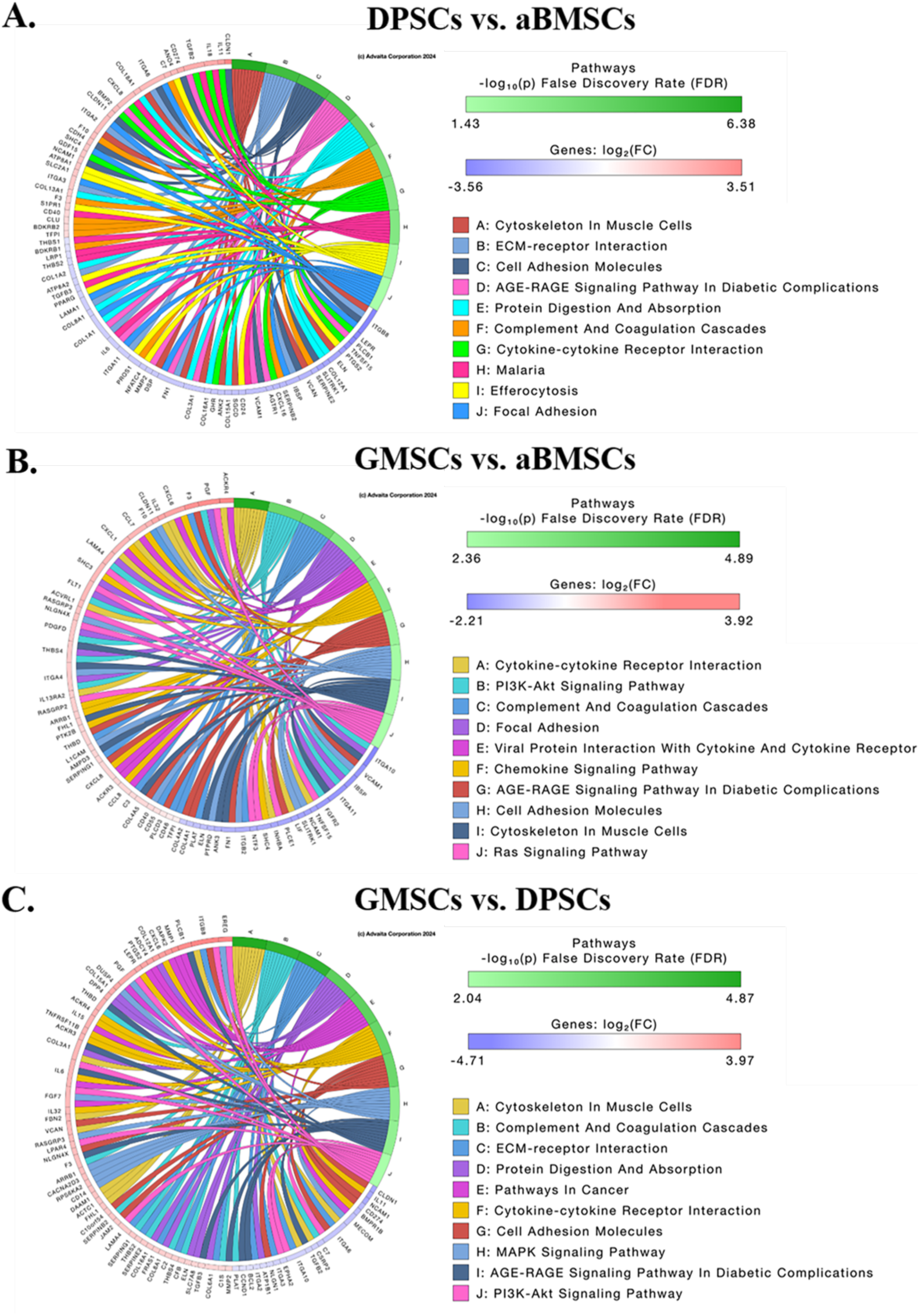
Chord diagram of Pairwise Comparisons. Chord Diagram of three different oral-derived MSC populations in a pairwise comparison showing the top 10 pathways and their corresponding 10 DE genes after p-value correction using FDR and log₂(FC), respectively. **A.** DPSCs vs. aBMSCs **B.** GMSCs vs. aBMSCs **C.** GMSCs vs. DPSCs. (osteoblast/chondrocyte/odontoblast differentiation) was upregulated, whereas VEGFA (endothelial differentiation) was downregulated in DPSCs vs. aBMSCs (Fig. S2A, Additional File 1). AXIN2 (odontoblast differentiation), FGF2 (Osteoblast/endothelial differentiation), SMAD1 (MSC differentiation), and NGF (neuron differentiation) were upregulated, whereas TEK (endothelial differentiation) was downregulated in GMSCs vs. DPSCs (Fig. S2B, Additional File 1). In GMSCs vs. aBMSCs, BDNF (neuron differentiation) was downregulated (Fig. S2C, Additional File 1). Heat maps of the top 50 DEGs and volcano plots of specific DEGs as explained above in pairwise comparisons were shown in Fig. 3B and Fig. 4A, respectively.

**Figure 6.**
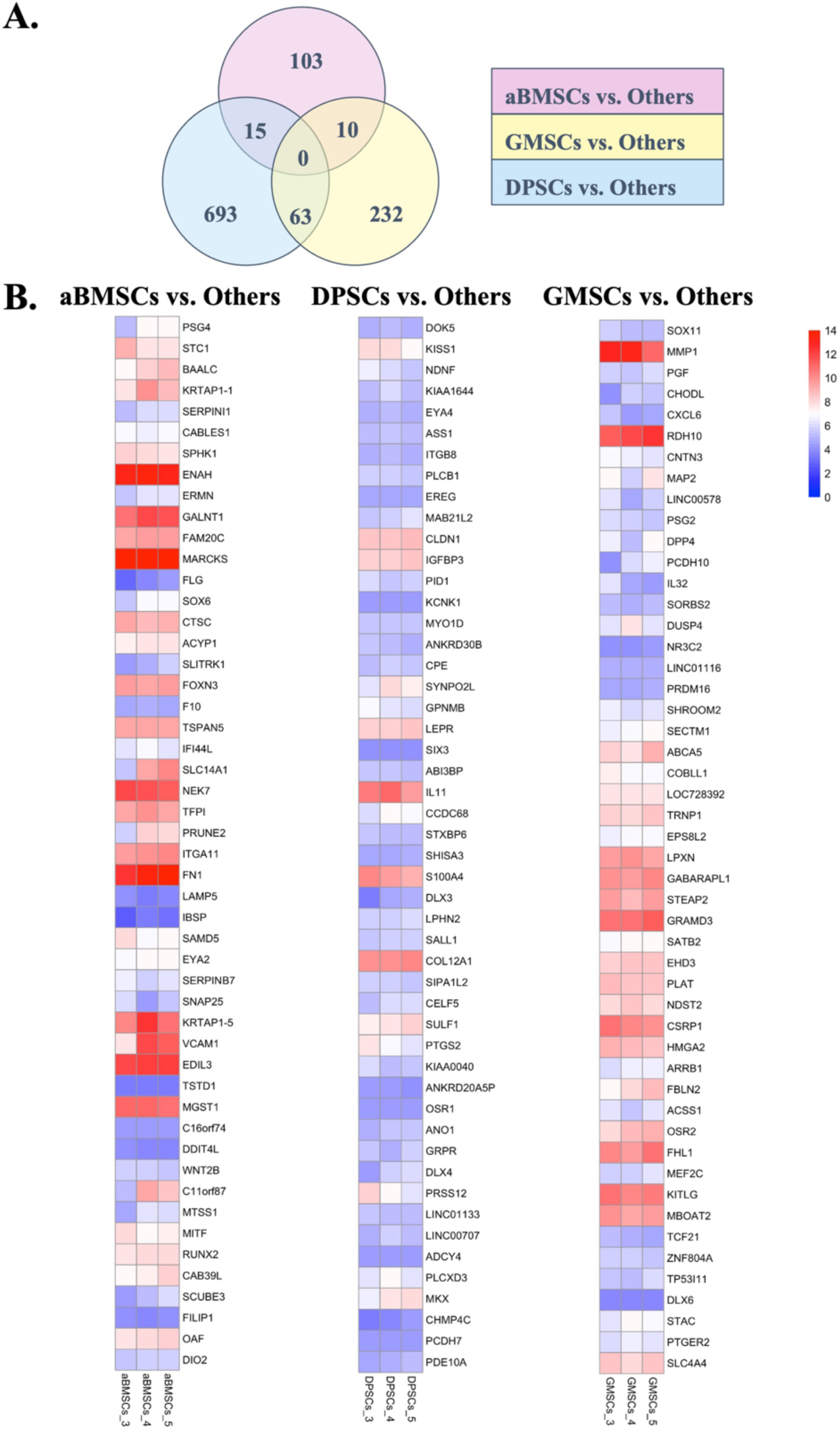
Venn Diagram and Heat Maps of Tissue-Specific Comparisons. **A.** Venn diagram obtained by metanalysis of all DEGs in aBMSCs vs. Others, DPSCs vs. Others, and GMSCs vs. Others. **B.** Heat maps of three different oral-derived MSC populations in a tissue-specific comparison showing the top 50 genes in aBMSCs vs. Others, DPSCs vs. Others, and GMSCs vs. Others according to their significance (FDR, *p*<0.05)

#### Pathways specific to the MSC population

The top 10 pathways with their corresponding genes for each tissue-specific comparison, aBMSCs vs. Others, DPSCs vs. Others, and GMSCs vs. Others after p-value correction with FDR (*p<0.05*) were shown in Fig. 7. Interestingly, after a meta-analysis of these comparisons, it was found that there was no unique pathway in aBMSCs vs. Others, four pathways were unique in DPSCs vs. Others. In contrast, the highest number of unique pathways (18 in total) were present in GMSCs vs. Others (Table S5, Additional File 1).

**Figure 7.**
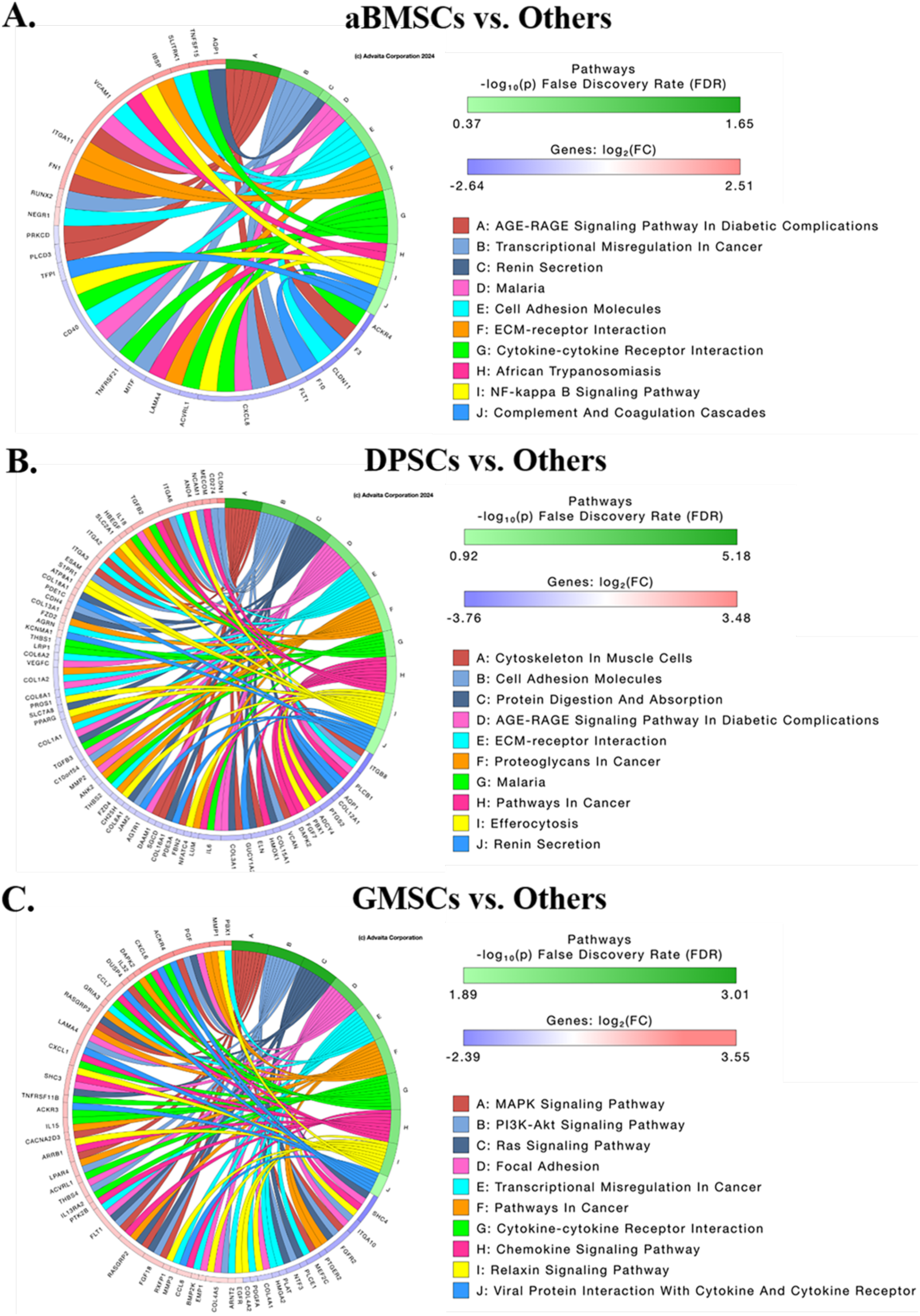
Chord diagram of Tissue-Specific Comparisons. Chord Diagram of three different oral-derived MSC populations in a tissue-specific comparison showing the top 10 pathways and their corresponding 10 DE genes after p-value correction using FDR and log₂(FC), respectively. **A.** aBMSCs vs. others **B.** DPSCs vs. others **C.** GMSCs vs. others.

#### Biological processes

It was found that only 896/26,994 GOBPs were found to be statistically significant (FDR, *p*<0.05) in all tissue-specific comparisons. Of these 896 GOBPs, 12/21 preselected biological processes were found with five biological processes unique to DPSCs (GO:0045669, GO:0045444, GO:0045599, GO:0032331, and GO:0016525), one GOBP unique to GMSCs (GO:0045668), and two GOBPs common to all MSCs (GO:0001525 and GO:0002062). Interestingly, no unique GOBP was identified in aBMSCs vs. Others, whereas some GOBPs are shared between two tissue-specific MSC populations. For example, aBMSCs and GMSCs share two GOBPs ([GO:0045766](https://www.ebi.ac.uk/QuickGO/term/GO:0002062) and GO:0032332). Similarly, GMSCs and DPSCs share two GOBPs (GO:0030182 and [GO:0001649](https://www.ebi.ac.uk/QuickGO/term/GO:0030182)), whereas DPSCs and aBMSCs share no GOBPs (Table 2).

**Table 2:**
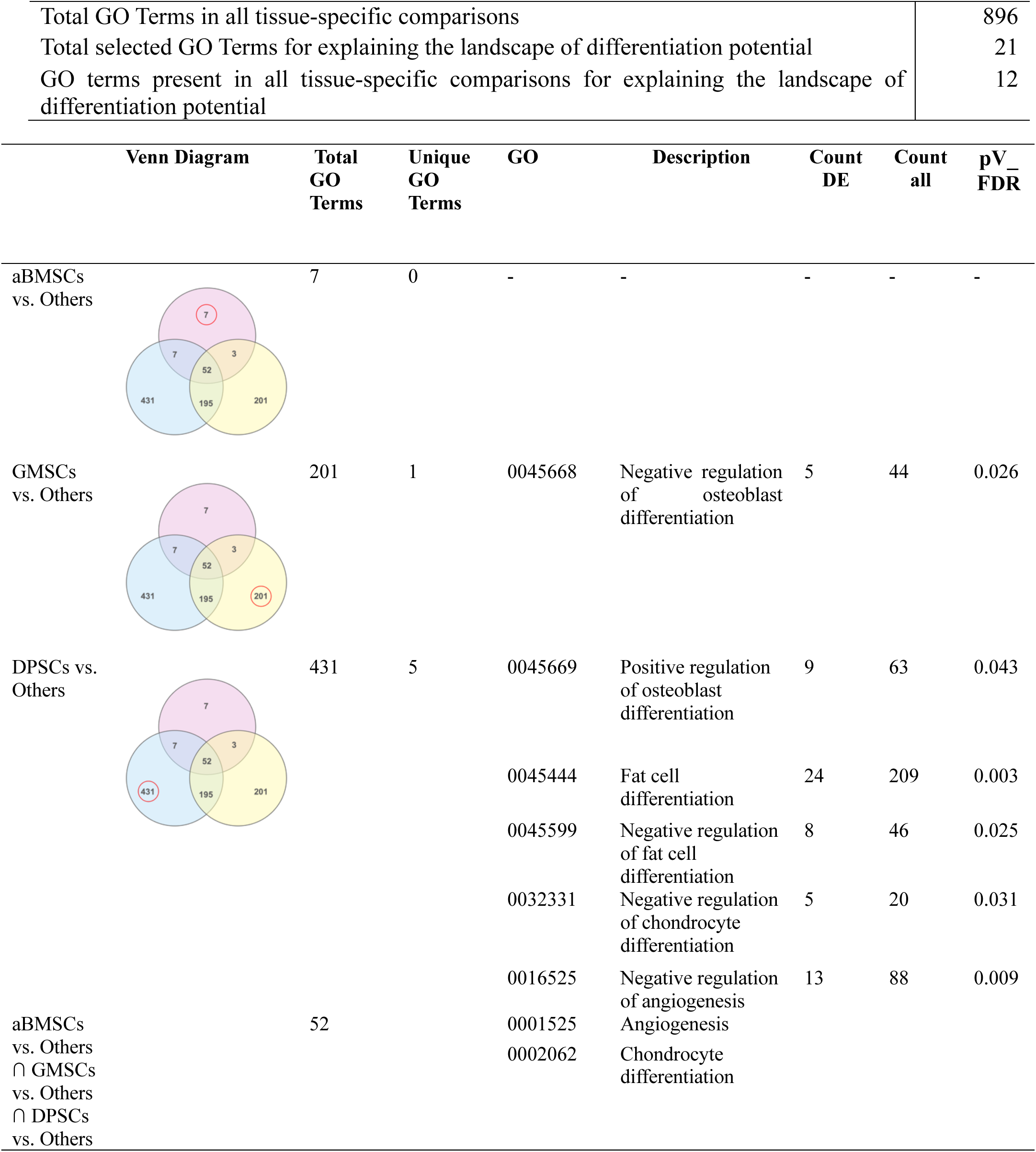
Tissue-specific comparison of three MSC populations showing unique GO terms, their DEseq genes, and the significance of these GO Terms uniquely to their populations (p-value correction: FDR.

| Total GO Terms in all tissue-specific comparisons |  |  |  |  |  |  | 896 |
| --- | --- | --- | --- | --- | --- | --- | --- |
| Total selected GO Terms for explaining the landscape of differentiation potential |  |  |  |  |  |  | 21 |
| GO terms present in all tissue-specific comparisons for explaining the landscape of differentiation potential |  |  |  |  |  |  | 12 |
| Venn Diagram | Total GO Terms | Unique GO Terms | GO | Description | Count DE | Count all | pV_FDR |
| aBMSCs vs. Others | 7 | 0 | - | - | - | - | - |
| GMSCs vs. Others | 201 | 1 | 0045668 | Negative regulation of osteoblast differentiation | 5 | 44 | 0.026 |
| DPSCs vs. Others | 431 | 5 | 0045669 | Positive regulation of osteoblast differentiation | 9 | 63 | 0.043 |
|  |  |  | 0045444 | Fat cell differentiation | 24 | 209 | 0.003 |
|  |  |  | 0045599 | Negative regulation of fat cell differentiation | 8 | 46 | 0.025 |
|  |  |  | 0032331 | Negative regulation of chondrocyte differentiation | 5 | 20 | 0.031 |
|  |  |  | 0016525 | Negative regulation of angiogenesis | 13 | 88 | 0.009 |
| aBMSCs vs. Others<br>∩ GMSCs vs. Others<br>∩ DPSCs vs. Others | 52 | | 0001525 | Angiogenesis | | | |
|  |  |  | 0002062 | Chondrocyte differentiation |  |  |  |

## Discussion

The increasing use of MSCs in regenerative medicine underscores the need for more rigorous characterization of these cells to ensure their efficacy and safety in clinical applications [30]. Through our established isolation and characterization techniques, we successfully delineated the distinct profiles of aBMSCs, DPSCs, and GMSCs. Leveraging bulk mRNA sequencing, we identified key DEGs in aBMSCs, DPSCs, and GMSCs with DPSC gene expression being the most distinct [31].

We compared the transcriptomic profiles of these different MSC populations from the same patients in a comprehensive attempt to better understand their intrinsic identities and differentiation potentials. The results highlighted the heterogeneity among these MSC populations, revealing distinct molecular signatures and pathways that drive their differentiation and immunomodulatory capacities in line with previous studies [31, 32].

Notably, our findings revealed significant differences in the expression of genes related to osteogenesis, neurogenesis, angiogenesis, and other crucial biological processes among the three oral-derived MSC populations. The osteogenic differentiation potential of aBMSCs, DPSCs, and GMSCs has been extensively reported, with both GMSCs and aBMSCs showing higher osteogenic capacity compared to DPSCs in different contexts [33, 34]. Our study supports these findings, indicating significant upregulation of osteogenic markers such as RUNX2 and IBSP in aBMSCs and SEMA4D in GMSCs but a downregulation of BMP4 in DPSCs. Even though GMSCs displayed a negative regulation of osteoblast differentiation in comparison with positive regulation of osteoblast differentiation in DPSCs, the unique set of pathways, such as the RAS1, RAP1, MAPK, and calcium signaling pathways, may contribute to their higher osteoblastic differentiation capabilities relative to DPSCs. However, relative to aBMSCs, GMSCs do not have this same innate osteogenic differentiation potential, as evidenced by the upregulation of specific osteoblastic genes such as RUNX2 and IBSP in aBMSCs.

The transcriptomic analysis also revealed DEGs involved in chondrogenesis and inflammation. SMAD7, SOX6, and ACAN were significantly upregulated in aBMSCs, suggesting their enhanced chondrogenic potential. This is consistent with prior studies highlighting the superior chondrogenic differentiation of bone-derived MSCs compared to other sources [7]. Conversely, IL6, a gene associated with inflammatory responses, was significantly downregulated in DPSCs, indicating their potential for reduced immunogenicity, which makes them a good candidate cell population for therapeutic clinical applications [8].

Our findings on the neurogenic differentiation potential are also consistent with prior studies that identified the expression of neurogenic markers in dental-derived MSCs. In GMSCs vs. DPSCs, NGF (neuron differentiation) was upregulated, whereas in GMSCs vs. aBMSCs, BDNF was downregulated. On further analyses, it was found that NES was significantly upregulated in DPSCs, whereas GDNF was significantly downregulated in GMSCs. Because of their neural crest origin, DPSCs express high mRNA expression of various neural crest developmental genes and neural crest-related genes [35], such as NES [36]. Our results aligned with a previous report where DPSCs exhibited the best neurogenic differentiation potential [37, 38]. However, in contrast, another study reported that GMSCs had higher neurogenic potential compared to DPSCs, stem cells from apical papilla (SCAPs), or MSCs from bone marrow (BMSCs) [39].

Our pathway analysis identified several pathways that were either unique to specific MSC populations or common across multiple comparisons. The PI3/AKT pathway, for instance, is well documented in the literature for promoting odontogenic [40] and osteogenic differentiation [41]. In contrast, in our study, we did not find the PI3/AKT pathway common in all MSC populations except GMSCs, plausibly due to our predefined criteria of significance cutoff of FDR <0.05. Other cell-cell interaction pathways, such as Cytokine-cytokine receptor interaction, ECM-receptor interaction, and Cell adhesion molecules, are statistically significant in all three populations of MSCs, underlying the important role of cell-cell interactions in various pathophysiological processes.

Despite the significant findings, our study has several limitations. Firstly, the sample size was limited to tissues from three patients, which may not fully represent the broader population’s genetic and phenotypic diversity. Future studies should include a larger sample size and a more diverse population (age, gender, race) to validate and uncover additional insights into MSC heterogeneity. Secondly, using bulk RNA sequencing, while powerful, may mask heterogeneity within individual cell populations. Single-cell RNA sequencing would provide a more robust understanding of cellular diversity and trajectories and identify specific subpopulations with distinct characteristics revealing potential therapeutic avenues. Thirdly, the study focused on MSCs derived from specific dental tissues while there are other sources of MSCs with different characteristics that were not explored. Therefore, expanding the sources of MSCs in future studies by including other dental tissues will provide a more comprehensive understanding of MSC heterogeneity, their characteristics, and potential applications. As a result, comparative studies across various MSC sources can help identify the most suitable types for specific therapeutic purposes. Finally, our functional assays, while demonstrating differentiation potential, were limited to *in vitro* conditions. Evaluating these cell populations in different in *vivo* contexts will be impactful in determining the therapeutic potential and safety of MSCs in clinical applications.

Our study highlights the intricate interplay between the tissue of origin and the microenvironment in shaping the “differentiation landscape” of oral-derived MSCs. Understanding the signaling pathways and genetic factors that drive MSC behavior will enable better design of therapeutic interventions that optimize their regenerative capabilities. Therefore, MSCs hold immense promise in the realm of regenerative medicine due to their capacity for tissue repair and regeneration

## Conclusion

In conclusion, our study confirms the diverse molecular landscapes of aBMSCs, DPSCs, and GMSCs, with DPSCs exhibiting the most unique molecular profile of the three. Furthermore, our findings lay the foundation for future longitudinal studies aimed at deciphering the key regulatory controls governing MSC differentiation, thereby advancing the field of regenerative medicine toward more efficacious and targeted therapeutic interventions.

## List of abbreviations

MSCs: Mesenchymal stem cells
aBMSCs: Alveolar bone-derived mesenchymal stem cells
DPSCs: Dental pulp-derived stem cells
GMSCs: Gingiva-derived mesenchymal stem cells
mRNA: Messenger ribonucleic Acid
DEGs: differentially expressed genes
FDR: False discovery rate
ISCT: International Society for Cellular Therapy
RNA-seq: RNA sequencing
IRB: Institutional Review Board
α MEM: α modification of minimum essential media
PBS: Phosphate-buffered saline
FBS: Fetal bovine serum
AA2P: L-ascorbic acid-2-phosphate
P: Passage
ARS: Alizarin Red S
HS: Histochemical Staining
AIM: Adipogenic Induction Media
AMM: Adipogenic Maintenance Media
ORO: Oil Red O
RT: Room temperature
PFA: Paraformaldehyde
BSA: Bovine serum albumin
PCR: Polymerase chain reaction
TMM: Trimmed mean of M
DE: Differentially expressed
DEGs: Differentially expressed genes
CD: Cluster of differentiation
SD: standard deviation
FDR: false discovery rate
SMAD1: SMAD family member 1
TWIST1: Twist family bHLH transcription factor 1
TWIST2: Twist family bHLH transcription factor 2
SOX6: SRY-box transcription factor 6
SOX9: SRY-box transcription factor 9
ALPL: Alkaline phosphatase, biomineralization associated
RUNX2: RUNX family transcription factor 2
IBSP. BSP 2: Integrin binding sialoprotein
BGLAP, Osteocalcin: Bone gamma-carboxyglutamate protein
BMP 2: Bone morphogenetic protein 2
BMP 4: Bone morphogenetic protein 4
BMP 9: GDF2 growth differentiation factor 2/Bone morphogenetic protein 9
COL1A1: Collagen type II alpha 1 chain
COL1A2: Collagen type II alpha 2 chain
MSX2: Msh homeobox 1
SHOX2: SHOX homeobox 2
PPARG: Peroxisome proliferator-activated receptor gamma
CEBPA: CCAAT enhancer binding protein alpha
ADIPOQ: Adiponectin, C1Q, and collagen domain-containing
AXL: AXL receptor tyrosine kinase
SMAD7: SMAD family member 7
BMP7: Bone morphogenetic protein 7
SOX9: SRY-box transcription factor 9
TGFβ1: Transforming growth factor beta 1
COL2A1: Collagen type II alpha 1 chain
ACAN: Aggrecan
NES: Nestin
SEMA3A: Semaphorin 3A
SEMA4D: Semaphorin 4D
BDNF: Brain-derived neurotrophic factor
GDNF: Glial cell-derived neurotrophic factor
NGF: Nerve growth factor
PGDFA: Platelet-derived growth factor subunit A
VEGFA: Vascular endothelial growth factor A
COL18A1: Collagen type XVIII alpha 1 chain
TEK: TEK receptor tyrosine kinase
ANGPT1: Angiopoietin 1
AXIN2: Axin 2
PAX9: Paired box 9
MSCs: Mesenchymal stem cells
aBMSCs: Alveolar bone-derived mesenchymal stem cells
DPSCs: Dental pulp-derived stem cells
GMSCs: Gingiva-derived mesenchymal stem cells
mRNA: Messenger ribonucleic Acid
FDR: False discovery rate
ISCT: International Society for Cellular Therapy
RNA-seq: RNA sequencing
IRB: Institutional Review Board
α MEM: α modification of minimum essential media
PBS: Phosphate-buffered saline
FBS: Fetal bovine serum
AA2P: L-ascorbic acid-2-phosphate
P: Passage
ARS: Alizarin Red S
HS: Histochemical Staining
AIM: Adipogenic Induction Media
AMM: Adipogenic Maintenance Media
ORO: Oil Red O
RT: Room temperature
PFA: Paraformaldehyde
BSA: Bovine serum albumin
PCR: Polymerase chain reaction
TMM: Trimmed mean of M
DE: Differentially expressed
DEGs: Differentially expressed genes
CD: Cluster of differentiation

## Declarations

### Ethics approval and consent to participate

This study was reviewed and approved by the University of Michigan IRB #HUM00142680. Informed consent was obtained from all the subjects who participated in the study.

### Consent for publication

Not applicable

### Availability of data and materials

Data associated with this study has not been deposited into a publicly available repository. However, all relevant data to support this study was included in the article and supplemental materials.

### Competing interests

The authors declare that they have no known competing financial interests.

### Funding

This project was supported with funding from the National Institutes of Health, and the National Institute of Dental and Craniofacial Research [R01DE028657 to DK].

### Authors’ contribution

HC: Writing – original draft, Writing – review & editing, Visualization, Validation, Methodology, Formal analysis, Investigation, Data curation, Conceptualization. CC: Writing – review & editing, Investigation, Visualization, Formal analysis, Data curation, Conceptualization. AH: Formal analysis and Data curation. SK: Formal analysis and Data curation. BM: Investigation and Methodology. RT: Writing – review & editing, Visualization, Investigation, Formal analysis. LG: Writing – review & editing, Visualization, Investigation, Formal analysis. JS-Writing – review & editing, Project administration. DK: Writing – review & editing, Supervision, Resources, Project administration, Investigation, Funding acquisition, Data curation, Conceptualization.

## Acknowledgments

The authors would like to acknowledge Kamlai Saiya cork and staff at the University of Michigan BSRB Flow Cytometry Core, Olivia Koues and staff at the University of Michigan Bioinformatics Core, and Melissa Coon, at the Advanced genomic core.

## Authors’ information

Department of Periodontics and Oral Medicine, School of Dentistry, University of Michigan, 1011 N. University, Ann Arbor, MI-48109, USA.

Hitesh Chopra, Chen Cao, Alice Hermann, Sumukh Kak, Jim Sugai & Darnell Kaigler

Department of Biomedical Engineering, College of Engineering, University of Michigan, 1011 N. University, Ann Arbor, MI-48109, USA.

Darnell Kaigler

Bioinformatics Core, Michigan Medicine, University of Michigan, North Campus Research Complex (NCRC), Building 22, 2800 Plymouth Road, Ann Arbor, MI-48109, USA.

Rebecca Tagett

Computational Medicine and Bioinformatics, Michigan Medicine, University of Michigan, North Campus Research Complex (NCRC), Building 22, Ann Arbor, MI-48109, USA.

Lana Garmire

## Notes

### Competing Interest Statement

The authors have declared no competing interest.

