## Additional File 1 for "Landscape of Differentiation Potentials as a “Hallmark” in Oral-derived MSCs"

Darnell Kaigler, D.D.S., M.S., Ph.D.

Associate Professor of Dentistry

Department of Periodontics and Oral Medicine

School of Dentistry, University of Michigan,

Ann Arbor, MI-48109, USA

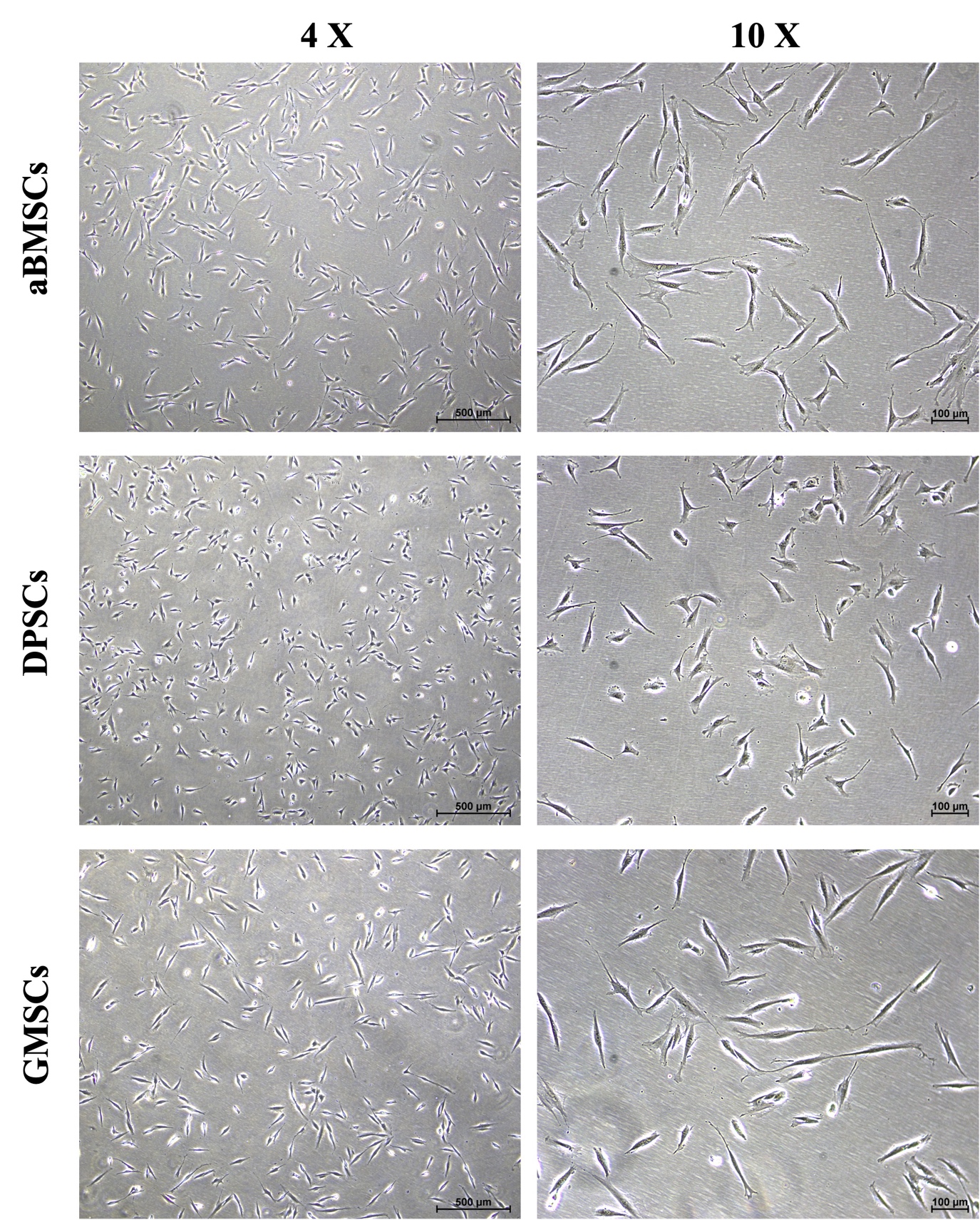

**Figure S1. Representative microscopic examination of the isolated aBMSCs, DPSCs, and GMSCs at 4 X and 10 X.**

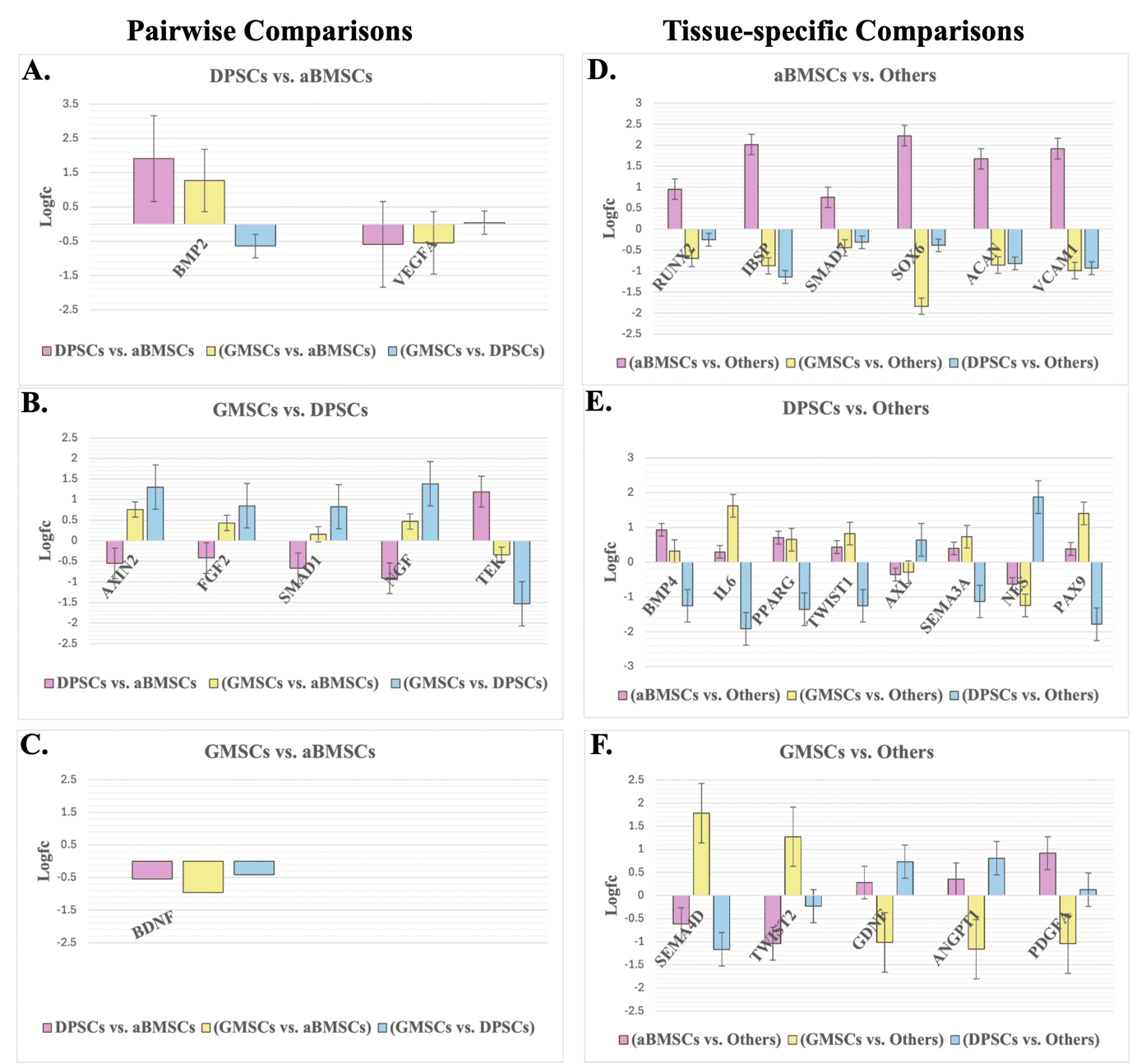

**Figure S2. Bar graphs with Standard error showing regulation of selected DEGs in Pairwise comparison (A-C) and Tissue-specific comparison (D-F).**

**Table S1: Flow Cytometer antibodies**

| ***Antibody** | **Biolegend Catalogue number** | **Clone** | **Conjugated w/ Fluorochrome** | **Ex. Laser** |
| --- | --- | --- | --- | --- |
| **CD 73** | 344004 | AD2 | PE | Yellow-Green Laser (561 nm) |
| **CD 90** | 328108 | 5E10 | FITC | Blue Laser (488 nm) |
| **CD 90 IC** | 400110 | MOPC-21 | FITC | Blue Laser (488 nm) |
| **CD 105** | 323206 | 43A3 | PE | Yellow-Green Laser (561 nm) |
| **CD 105 IC** | 400114 | MOPC-21 | PE | Yellow-Green Laser (561 nm) |
| **CD 34** | 343516 | 581 | PE/Cy7 | Yellow-Green Laser (561 nm) |
| **CD34 IC** | 400126 | MOPC-21 | PE/Cy7 | Yellow-Green Laser (561 nm) |
| **CD 45** | 304024 | HI30 | Alexa Fluor® 700 | Red Laser (633 nm) |
| **CD 45 IC** | 400144 | MOPC-21 | Alexa Fluor® 700 | Red Laser (633 nm) |

***All antibodies are specific to humans with Mouse IgG1, κ isotype control**

**Table S2: Notable genes to explain the landscape of differentiation potential in all three MSC populations**

|  | **Gene** | **Gene ID** |
| --- | --- | --- |
|  | **Mesenchymal stem cell differentiation** | |
| 1. | SMAD family member 1 (SMAD1) | [4086](https://www.ncbi.nlm.nih.gov/gene/4086) |
| 2. | Twist family bHLH transcription factor 1 (TWIST1) | [7291](https://www.ncbi.nlm.nih.gov/gene/7291) |
| 3. | Twist family bHLH transcription factor 2 (TWIST2) | [117581](https://www.ncbi.nlm.nih.gov/gene/?term=117581) |
| 4. | SRY-box transcription factor 6 (SOX6) | [55553](https://www.ncbi.nlm.nih.gov/gene/55553) |
| 5. | SRY-box transcription factor 9 (SOX9) | [6662](https://www.ncbi.nlm.nih.gov/gene/6662) |
|  | **Osteogenic Induction/differentiation** | |
| 6. | Alkaline phosphatase, biomineralization associated (ALPL) | [249](https://www.ncbi.nlm.nih.gov/gene?Db=gene&Cmd=DetailsSearch&Term=249) |
| 7. | RUNX family transcription factor 2 (RUNX2) | [860](https://www.ncbi.nlm.nih.gov/gene/860) |
| 8. | Integrin binding sialoprotein (IBSP. BSP 2) | [3381](https://www.ncbi.nlm.nih.gov/gene/?term=3381%5Buid%5D) |
| 9. | Bone gamma-carboxyglutamate protein (BGLAP, Osteocalcin) | [632](https://www.ncbi.nlm.nih.gov/gene/?term=632%5Buid%5D) |
| 10. | Bone morphogenetic protein 2 (BMP 2) | [650](https://www.ncbi.nlm.nih.gov/gene/?term=650%5Buid%5D) |
| 11. | Bone morphogenetic protein 4 (BMP 4) | [652](https://www.ncbi.nlm.nih.gov/gene/652) |
| 12. | GDF2 growth differentiation factor 2/Bone morphogenetic protein 9 (BMP 9) | [2658](https://www.ncbi.nlm.nih.gov/gene/?term=2658%5Buid%5D) |
| 13. | Collagen type II alpha 1 chain (COL1A1) | [1277](https://www.ncbi.nlm.nih.gov/gene/1277) |
| 14. | Collagen type II alpha 2 chain (COL1A2) | [1288](https://www.ncbi.nlm.nih.gov/gene/1278) |
| 15. | Msh homeobox 1 (MSX2) | [4488](https://www.ncbi.nlm.nih.gov/gene/4488) |
| 16. | SHOX homeobox 2 (SHOX2) | [6474](https://www.ncbi.nlm.nih.gov/gene/6474) |
|  | **Adipogenic Induction/differentiation** | |
| 17.  | Peroxisome proliferator activated receptor gamma (PPARG) | [5468](https://www.ncbi.nlm.nih.gov/gene/?term=5468%5Buid%5D) |
| 18.  | CCAAT enhancer binding protein alpha (CEBPA) | [1050](https://www.ncbi.nlm.nih.gov/gene/1050) |
| 19.  | Adiponectin, C1Q and collagen domain containing (ADIPOQ) | [9370](https://www.ncbi.nlm.nih.gov/gene/?term=9370%5Buid%5D) |
|  | Chondrogenic Induction/differentiation |  |
| 20.  | AXL receptor tyrosine kinase (AXL) | [558](https://www.ncbi.nlm.nih.gov/gene/558) |
| 21.  | SMAD family member 7 (SMAD7) | <4092> |
| 22.  | Bone morphogenetic protein 2 (BMP2) | [650](https://www.ncbi.nlm.nih.gov/gene/?term=650%5Buid%5D) |
| 23.  | Bone morphogenetic protein 4 (BMP4) | [652](https://www.ncbi.nlm.nih.gov/gene/652) |
| 24.  | Bone morphogenetic protein 7 (BMP7) | [655](https://www.ncbi.nlm.nih.gov/gene/655) |
| 25.  | SRY-box transcription factor 9 (SOX9) | [6662](https://www.ncbi.nlm.nih.gov/gene/6662) |
| 26.  | Transforming growth factor beta 1 (TGFβ1) | [7040](https://www.ncbi.nlm.nih.gov/gene/7040) |
| 27.  | Collagen type II alpha 1 chain (COL2A1) | [1280](https://www.ncbi.nlm.nih.gov/gene/1280) |
| 28.  | Aggrecan (ACAN) | [176](https://www.ncbi.nlm.nih.gov/gene/176) |
|  | **Neurogenic Induction/differentiation** | |
| 29.  | Nestin (NES) | [10763](https://www.ncbi.nlm.nih.gov/gene/10763) |
| 30.  | Semaphorin 3A (SEMA3A) | [10371](https://www.ncbi.nlm.nih.gov/gene/?term=10371) |
| 31.  | Semaphorin 4D (SEMA4D) | [10507](https://www.ncbi.nlm.nih.gov/gene/?term=10507%5Buid%5D) |
| 32.  | Bone morphogenetic protein 9 (BMP9) | [2658](https://www.ncbi.nlm.nih.gov/gene/2658) |
| 33.  | Transforming growth factor beta 1 (TGFβ1) | [7040](https://www.ncbi.nlm.nih.gov/gene/7040) |
| 34.  | Brain derived neurotrophic factor (BDNF) | [627](https://www.ncbi.nlm.nih.gov/gene/627) |
| 35.  | Glial cell derived neurotrophic factor (GDNF) | [2668](https://www.ncbi.nlm.nih.gov/gene/2668) |
| 36.  | Nerve growth factor (NGF) | [4803](https://www.ncbi.nlm.nih.gov/gene/4803) |
|  | Angiogenic induction/differentiation |  |
| 37.  | Platelet derived growth factor subunit A (PGDFA) | [5154](https://www.ncbi.nlm.nih.gov/gene/5154) |
| 38.  | Vascular endothelial growth factor A (VEGFA) | [7422](https://www.ncbi.nlm.nih.gov/gene/?term=7422%5Buid%5D) |
| 39.  | Collagen type XVIII alpha 1 chain (COL18A1) | [80781](https://www.ncbi.nlm.nih.gov/gene/?term=80781%5Buid%5D) |
| 40.  | TEK receptor tyrosine kinase (TEK) | [7010](https://www.ncbi.nlm.nih.gov/gene/?term=7010%5Buid%5D) |
| 41.  | Angiopoietin 1 (ANGPT1) | [284](https://www.ncbi.nlm.nih.gov/gene/284) |
|  | **Odontogenic induction/differentiation** | |
| 42.  | Axin 2 (AXIN2) | [8313](https://www.ncbi.nlm.nih.gov/gene/8313) |
| 43.  | Bone morphogenetic protein 2 (BMP2) | [650](https://www.ncbi.nlm.nih.gov/gene/?term=650%5Buid%5D) |
| 44.  | Paired box 9 (PAX9) | [5083](https://www.ncbi.nlm.nih.gov/gene/5083) |
|  | **Inflammatory** |  |
| 46 | VCAM1 | [7412](https://www.ncbi.nlm.nih.gov/gene/7412) |
| 47 | IL6 | [3569](https://www.ncbi.nlm.nih.gov/gene/7412) |

**Table S3: 21 Selected biological processes to explain the landscape of differentiation potential in all three MSC populations**

|  | **Cells** | **GO Term** |  |
| --- | --- | --- | --- |
| 1. | MSCs | Mesenchymal Stem Cell Differentiation | GO:0072497 |
|  |  | Positive Regulation Of Mesenchymal Stem Cell Differentiation | GO:2000741 |
|  |  | Negative Regulation Of Mesenchymal Stem Cell Differentiation | GO:2000740 |
| 2. | Osteoblasts | Osteoblast Differentiation | GO:0001649 |
|  |  | Positive Regulation Of Osteoblast Differentiation | GO:0045669 |
|  |  | Negative Regulation Of Osteoblast Differentiation | GO:0045668 |
| 3. | Adipocytes | Fat Cell Differentiation | GO:0045444 |
|  |  | Positive Regulation Of Fat Cell Differentiation | GO:0045600 |
|  |  | Negative Regulation Of Fat Cell Differentiation | GO:0045599 |
| 4. | Chondrocytes | Chondrocyte Differentiation | GO:0002062 |
|  |  | Positive Regulation Of Chondrocyte Differentiation | GO:0032332 |
|  |  | Negative Regulation Of Chondrocyte Differentiation | GO:0032331 |
| 5. | Neurons | Neuron Differentiation | GO:0030182 |
|  |  | Positive Regulation Of Neuron Differentiation | GO:0045666 |
|  |  | Negative Regulation Of Neuron Differentiation | GO:0045665 |
| 6. | Endothelial cells | Angiogenesis | GO:0001525 |
|  |  | Positive Regulation Of Angiogenesis | GO:0045766 |
|  |  | Negative Regulation Of Angiogenesis | GO:0016525 |
| 7. | Odontoblasts | Odontoblast Differentiation | GO:0071895 |
|  |  | Positive Regulation Of Odontoblast Differentiation | GO:1901331 |
|  |  | Negative Regulation Of Odontoblast Differentiation | GO:1901330 |

**Table S4: Selected biological pathways to explain the landscape of differentiation potential in pairwise comparisons**

|  | Venn diagram | Total # of pathways | Relevant pathway description | pV_FDR |
| --- | --- | --- | --- | --- |
| DPSCs vs aBMSCs | 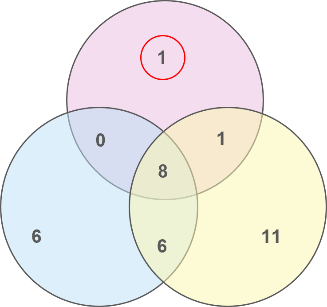 | 1 | - | - |
| GMSCs vs aBMSCs | 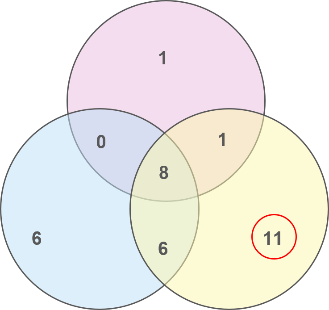 | 11 | Calcium signaling pathway | 0.013 |
|  |  |  | Chemokine signaling pathway | 8.891e-4 |
|  |  |  | IL-17 signaling pathway | 0.050 |
|  |  |  | RAP1 signaling pathway | 0.004 |
|  |  |  | RAS1 signaling pathway | 0.004 |
| GMSCs vs DPSCs | 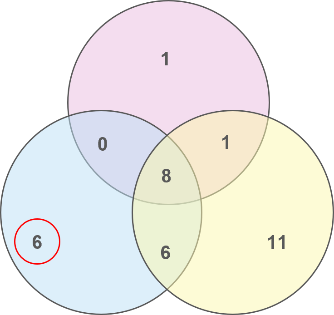 | 6 | TGF-ü signaling pathway | 0.009 |

| 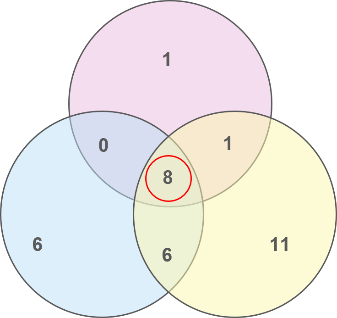 | **Pathways (8)** | pV_FDR (DPSCs vs aBMSCs) | pV_FDR (GMSCs vs aBMSCs) | pV_FDR (GMSCs vs DPSCs) |
| --- | --- | --- | --- | --- |
|  | Cell adhesion molecules | 0.00001 | 0.00356 | 0.00046 |
|  | Complement and coagulation cascades | 0.00066 | 0.00031 | 0.00003 |
|  | Cytokine-cytokine receptor interaction | 0.00125 | 0.00001 | 0.00039 |
|  | Cytoskeleton in muscle cells | 0.00000 | 0.00393 | 0.00001 |
|  | ECM-receptor interaction | 0.00001 | 0.00886 | 0.00003 |

**Table S5: Selected biological pathways to explain the landscape of differentiation potential in tissue-specific comparisons**

|  | Venn diagram | Total # of pathways | Relevant pathway description | pV_FDR |
| --- | --- | --- | --- | --- |
| aBMSCs vs others | 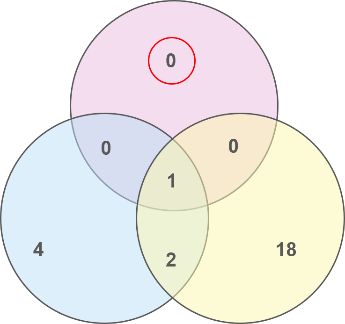 | 0 | - | - |
| GMSCs vs others | 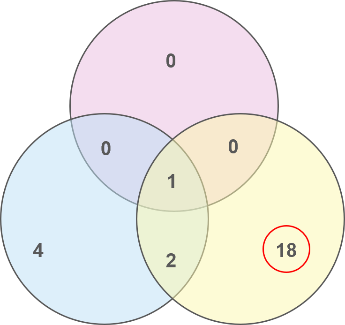 | 18 | Calcium signaling pathway | 0.033 |
|  |  |  | Chemokine signaling pathway | 0.006 |
|  |  |  | Complement and coagulation cascades | 0.036 |
|  |  |  | Cytokine-cytokine receptor interaction | 0.006 |
|  |  |  | IL-17 signaling pathway | 0.028 |
|  |  |  | MAPK signaling  pathway | 0.001 |
|  |  |  | PI3K-Akt signaling pathway | 0.001 |
|  |  |  | RAP1 signaling pathway | 0.014 |
|  |  |  | RAS1 signaling pathway | 0.001 |
| DPSCs vs others | 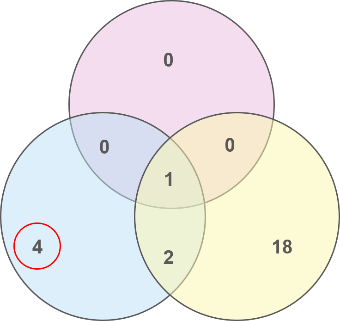 | 4 | Cell adhesion molecules | 0.0003 |
|  |  |  | ECM-receptor interaction | 0.0063 |
| aBMSCs vs Others ∩ GMSCs vs Others ∩ DPSCs vs Others | 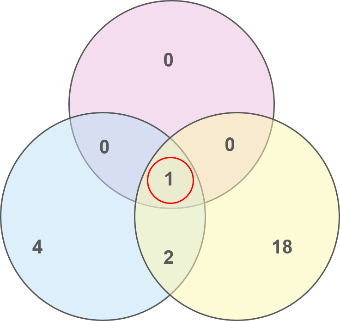 | 1 | - | - |
